# Baikal: Unpaired Denoising of Fluorescence Microscopy Images using Diffusion Models

**DOI:** 10.1101/2024.06.04.597486

**Authors:** Shivesh Chaudhary, Sivaramakrishnan Sankarapandian, Matt Sooknah, Joy Pai, Caroline McCue, Zhenghao Chen, Jun Xu

## Abstract

Fluorescence microscopy is an indispensable tool for biological discovery but image quality is constrained by desired spatial and temporal resolution, sample sensitivity, and other factors. Computational denoising methods can bypass imaging constraints and improve signal-to-noise ratio in images. However, current state of the art methods are commonly trained in a supervised manner, requiring paired noisy and clean images, limiting their application across diverse datasets. An alternative class of denoising models can be trained in a self-supervised manner, assuming independent noise across samples but are unable to generalize from available unpaired clean images. A method that can be trained without paired data and can use information from available unpaired highquality images would address both weaknesses. Here, we present Baikal, a first attempt to formulate such a framework using Denoising Diffusion Probabilistic Models (DDPM) for fluorescence microscopy images. We first train a DDPM backbone in an unconditional manner to learn generative priors over complex morphologies in microscopy images. We then apply various conditioning strategies to sample from the trained model and propose an optimal strategy to denoise the desired image. Extensive quantitative comparisons demonstrate better performance of Baikal over state of the art self-supervised methods across multiple datasets. We highlight the advantage of generative priors learnt by DDPMs in denoising complex *Flywing* morphologies where other methods fail. Overall, our DDPM based denoising framework presents a new class of denoising methods for fluorescence microscopy datasets that achieve good performance without collection of paired high-quality images. Github repo: https://github.com/scelesticsiva/denoising/tree/main

## 1 Introduction

Fluorescence microscopy is a widely used technique to study biological phenomena. Advancements in optical techniques [7] and development of better fluorescent sensors [46] have pushed the limits of image collection. However desired imaging conditions like spatio-temporal resolution, duration for time-lapse imaging, phototoxicity and photobleaching for sensitive samples etc. constrain imaging parameters such as laser power, exposure time and frame-rate leading to acquisition of noisy images [30, 37].

Several computational denoising methods have been developed for both single images [1, 4, 5, 8, 12, 15, 16, 18, 20, 25–28, 31–34, 39, 40, 42, 44, 47] and videos [6, 10, 17, 22] that can bypass imaging constraints and generate clean images. These methods fall into one of the two categories: 1) *paired-denoising* methods like CARE [42] offer best denoising performance, however these methods require large paired datasets of noisy and high quality images for training that are extremely time-consuming to collect. 2) *self-supervised denoising* methods such as Noise2Noise (N2N) [18], Noise2Void (N2V) [15], Noise2Self [1] etc. can be trained using only noisy images. However, these methods assume pixel-wise independent noise, achieving worse performance in situations where spatially correlated (structured) noise is present as shown in [3]. To account for structured noise, modifications of self-supervised denoising methods have been proposed such as StructN2V [3], SSID [19] etc. Here, we consider a common scenario where an unpaired dataset of clean images is available but not paired noisy-clean data. Unpaired clean images may provide important morphological priors to denoising methods in extreme noise scenarios. Previous methods are not designed to take advantage of such unpaired images.

Generative Adversarial Network (GAN) based approaches have been proposed for fluorescence microscopy image restoration using unpaired datasets [21, 29]. Howvwer GAN models suffer from instability during training thus leading to lower adoption among community. Denoising Diffusion Probabilistic Models (DDPMs) [13] have recently demonstrated superior performance over GANs in learning generative distribution of natural images and high quality image-synthesis [9, 35, 36]. But suitability and applicability of DDPMs for fluorescence microscopy denoising task has not been shown yet.

Our method, which we term Baikal, leverages a Denoising Diffusion Probabilistic Model (DDPM) [13] for learning generative priors from unpaired clean images. DDPMs have been applied for denoising data from several modalities such as PET, MRI, CT [11, 23, 38, 43]. However fluorescence microscopy data generation and noise characteristics differ significantly from these modalities [2]. Thus it is not clear whether DDPMs would work for denoising fluorescence microscopy images. We explore this question in this work by leveraging DDPM in a two step framework. First a diffusion model is trained only on clean images. Next, conditional sampling is used to generate clean predictions of the noisy images. In this work, we empirically evaluate several conditional samplers on multiple open-source datasets [42]. The proposed framework has several advantages over previous methods 1) it does not require paired data as *paired-denoising* methods for training, 2) the DDPM backbone learns important morphological priors from unpaired clean images to guide denoising, 3) it is widely applicable across datasets with varying noise properties. Our contributions are as follows,

**–** We demonstrate the first application of Diffusion Models for denoising fluorescence microscopy images.
**–** We systematically evaluate the applicability of several conditional sampling strategies designed for inpainting tasks for denoising and suggest an optimal strategy for fluorescence microscopy datasets.
**–** We evaluate whether outputs from self-supervised methods can better guide conditional samplers.

## 2 Methods

Our goal is to generate a clean image ***x*** given a noisy fluorescence image ***y*** . *Self-supervised* methods only use to generate clean images. *Paired-denoising* methods use paired clean and noisy acquisitions, to generate clean images. Here, we propose a two step method for denoising flourescence microscopy images when unpaired clean and noisy images are available (Fig. 1). First a generative backbone is trained to model clean data. Next, adapting several samplers from the inpainting literature [24] we evaluate them on three datasets of varying morphological properties. In addition, if *self-supervised* predictions are available for noisy images, we evaluate their utility on the performance of samplers.

**Fig. 1.**
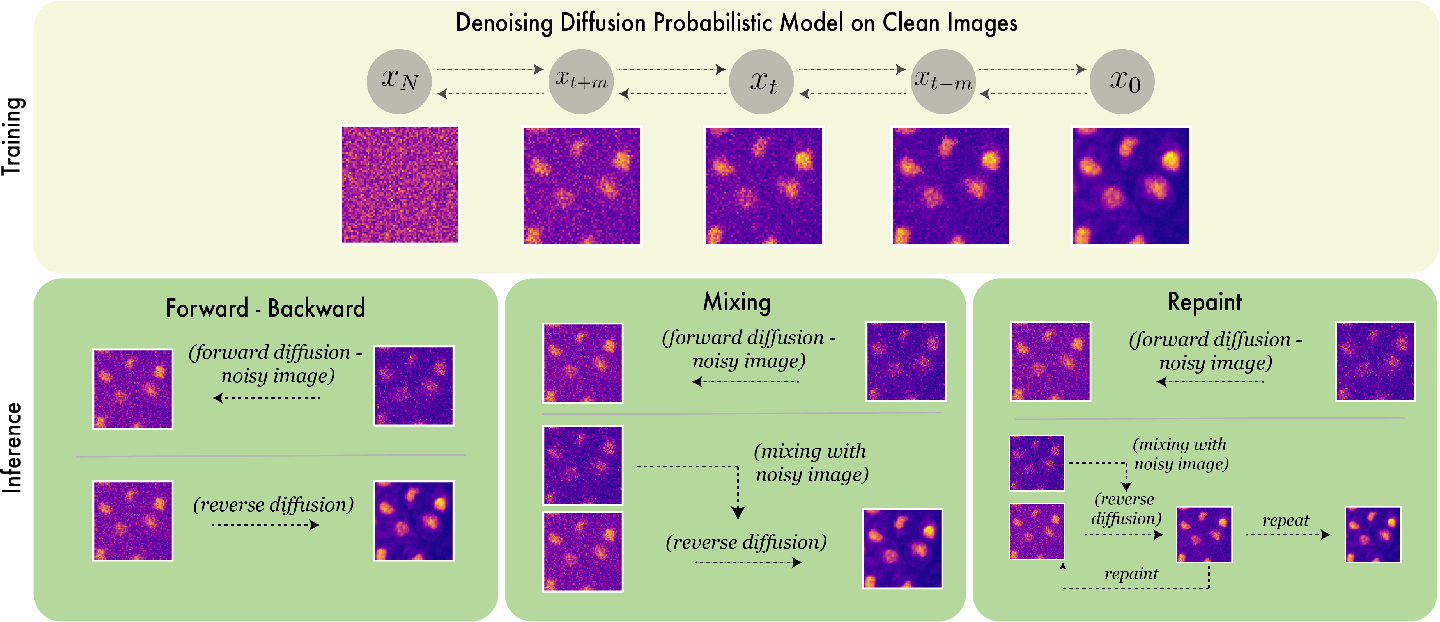
Overview of the proposed method. **Training stage** a DDPM backbone is trained using only unpaired high quality images. **Inference Stage** image is denoised by sampling from the trained backbone. Three strategies are explored to guide the sampling process namely Forward-Backward, Mixing and Repaint.

### 2.1 Generative backbone on clean images

Our generative backbone is trained as a DDPM [13] on unpaired clean images . DDPMs are generative latent variable models that learn a model distribution *p_θ_*(***x***) that approximate data distribution *q*(***x***). The *forward* diffusion process applies Gaussian noise to the images until they become indistinguishable from random Gaussian noise for *T* steps ***x***_0_ *→* ***x***_1_ *→* ***x***_2_*, …,* ***x****_T_* . Variance schedule is given by *β*_1_*, …, β_T_* .

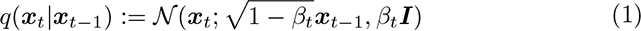

The *reverse* process goes in the opposite direction to generate images from random noise ***x****_T_ →* ***x****_T−_*_1_ *→* ***x****_T−_*_2_*, …,* ***x***_0_ where ***x****_T_ ∼ N* (**0**, ***I***). ***µ****_θ_* denotes the neural network trained to denoise ***x****_t_* at every time step *t*. Variances of the reverse process are also learned unless otherwise specified.

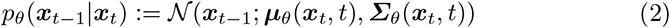

### 2.2 Conditional samplers as denoisers

Once the generative backbone is trained on clean images, we denoise an image ***y*_0_** by generating a (predicted) clean image ***x*_0_** conditioned on ***y*_0_**. We adapt conditional sampling methods proposed in the inpainting literature [24] and evaluate their performance in the context of denoising fluorescence images. All conditional samplers in this work first forward diffuse the noisy image ***y***_0_ to arbitrary time step *t* utilizing the closed form expression for forward process as follows,

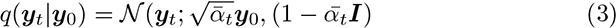

where 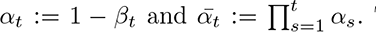 To evaluate N2V predictions for their utility in our proposed denoising method, we replace ***y***_0_ with ***y****^N^*^2*V*^ and its corresponding forward diffused as ***y****^N^*^2*V*^ . Once noisy images ***y***_0_ are forward diffused, different conditional sampling methods differ in the inputs to ***µ****_θ_*(***x****_t_, t*).

### Forward-Backward (FB)

Here, first the noisy image ***y*_0_** is forward diffused to *t^′^* steps such that only the underlying signal is preserved. Next, the reverse process is run by setting ***x****_t_′* = ***y****_t_′* (only at *t* = *t^′^*) to generate denoised ***x***_0_ from noisy ***y***_0_. The reverse step is given as follows,

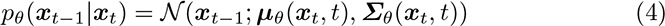

Intuitively, since the DDPM is trained only on clean images, at the end of the reverse process, our hope is that the generated sample will approximate the denoised image (i.e. ***x***_0_ ***x***). However one disadvantage of this sampler is that the original signal destroyed during forward diffusion is lost forever and the network might not be able to recover all the components of the clean image when running the reverse process. To run on N2V predictions, we replace ***x****_t_′* = ***y****^N^*^2*V*^ .

### Mixing

In forward-backward sampler the generation process at each time step (except for *t* = *t^′^*) is independent of the noisy image ***y*_0_** (i.e.) each generative step is dependent only on the previously generated image. Additionally, information once destroyed is not recoverable in forward-backward denoising. Thus to feed the network all the information that was present in the original noisy image at every time step and to further guide the generation process, we compute the weighted average of the reverse sample ***x****_t_* and the forward diffused noisy image ***y****_t_* and treat them as inputs to ***µ****_θ_*.

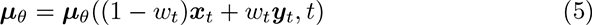

Since higher weights for the noisy image in the later stages of the generative process could yield noisy generations, we monotonically decrease the weights *w_t_* during the reverse process (i.e) *w_t_ > w_t−_*_1_. To run on N2V predictions, we replace 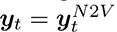 when taking the weighted combination, but use forward noised ***y***_0_ as ***y****_t_* only for *t* = *t^′^*. A disadvantage of this sampler is that, as we re-introduce noisy image at every time step by taking weighted combination, it replaces denoised pixels by noisy pixels reverting denoising.

### Repaint

Although weighted combinations of ***y****_t_* and DDPM outputs at each time step ***x****_t_* guide the generation process, some of the denoised pixels in ***x****_t_* are replaced with noisy pixels, thus reverting the denoising process. Additionally, mixing once at each time step may not be enough to ensure semantic consistency. This is because the mixing input ***y****_t_* at each time step is independent of the DDPM output ***x****_t_* at that time step. Thus, although the DDPM tries to predict the denoised output at the next time step using both ***y****_t_* and ***x****_t_*, it may not be able to correct any inconsistency in the previous step. Thus, to ensure semantic consistency, similar to Repaint [24], we repeat the reverse step multiple times. Since ***µ****_θ_* is trained as a *denoiser*, we use it for maintaining semantic consistency. Concretely, starting at a time step *t*, we run the the generation process for *U* steps, while mixing ***x****_t_* and ***y****_t_* at each time step, to generate ***x****_t−U_* . Next, ***x****_t−U_* is taken as input to the time step *t* and generation process is repeated again several times (See Algorithm 1). Our strategy is different from the original Repaint, in that, empirically we found repainting every step yielded worse results compared to repainting in intervals. Additionally, we stop mixing in the later stages of generation to avoid replacing denoised pixels by noisy pixels resulting in better results as shown in Fig.4(right).

## 3 Experiments

### 3.1 Dataset & Training

We evaluate our framework on three open source fluorescence microscopy datasets [42] with distinct morphological labelling to highlight the generalizability of Baikal. These include 1) nuclei labeled embryos of *Tribolium castaneum*, 2) nuclei labeled embryos of the planarian *S. mediterranea* and 3) boundary labelled eplithelia of the fruit fly *Drosophila Melanogaster*. The datasets provide noisy and high quality images that were acquired at low and high laser powers, respectively. The test sets provided in the datasets were not normalized as the train set. Thus, for fair evaluation, we split the provided train sets into train, eval and test sets in the ratio 80 : 10 : 10. We provide all qualitative and quantitative results on test splits. We train diffusion backbone using the DifFace [45] codebase^*^ on clean images from the train set.

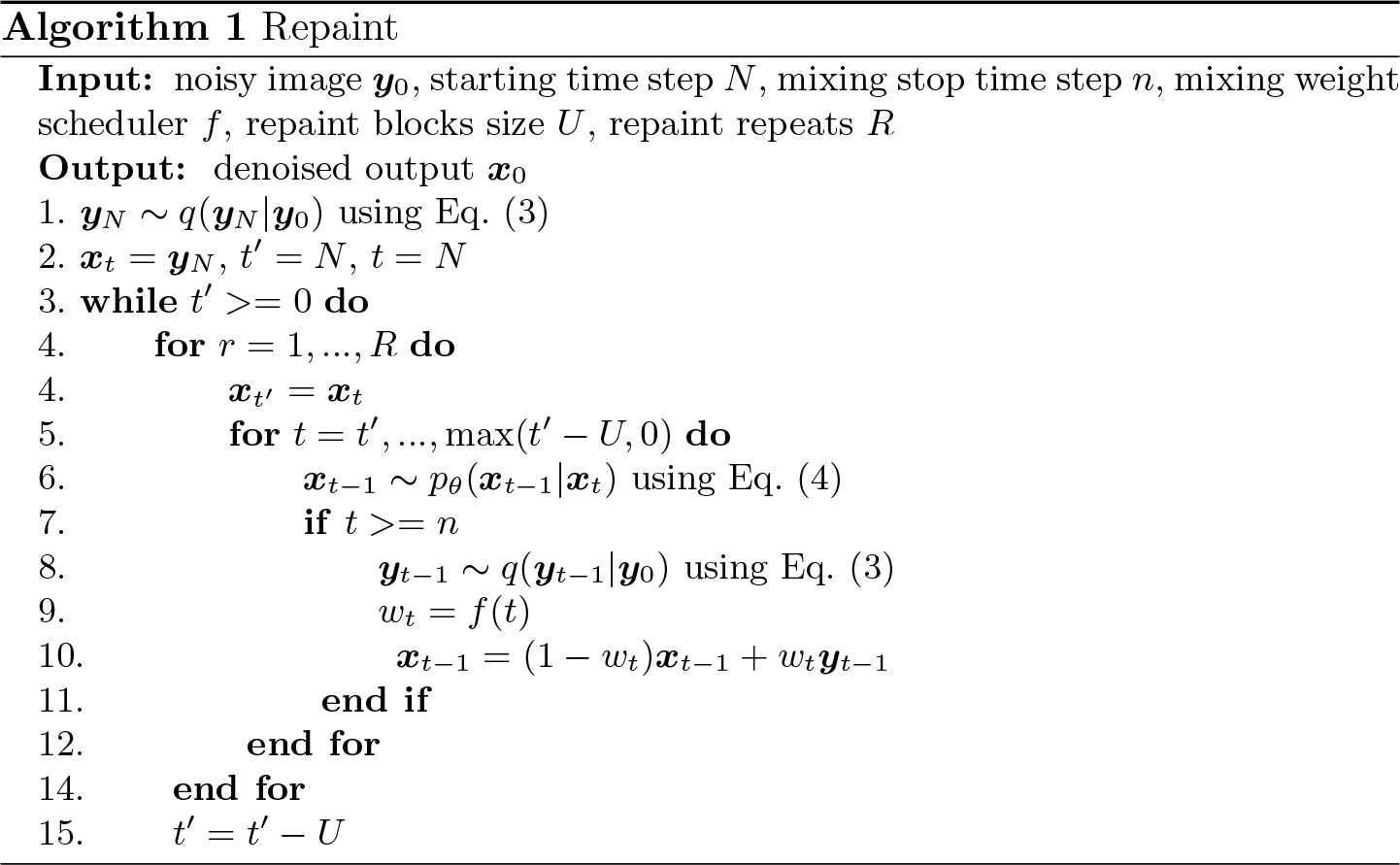

*Planaria* and *Tribolium* datasets include noisy and clean 3D volumes, thus for these datasets we train the diffusion backbone treating each 2D z-plane as individual image. Random z-plane images sampled from trained diffusion model for these datasets are shown in Fig.2. During inference, each individual z-plane is denoised independently by the trained diffusion model for these datasets. In comparison, *Drosophila* dataset consisted of 3D noisy images and 2D max-projected clean images. Thus for *Drosophila* dataset, we trained diffusion model on max-projected 2D clean images. Random max-projected images sampled from trained diffusion model for *Drosophila* dataset are shown in Fig.2. During inference on *Drosophila* dataset, each z-plane is denoised individually using the trained model. Subsequently, denoised 3D stack is max-projected for metrics evaluation. Thus, our framework does not require training a separate projector network as done previously [42].

**Fig. 2.**
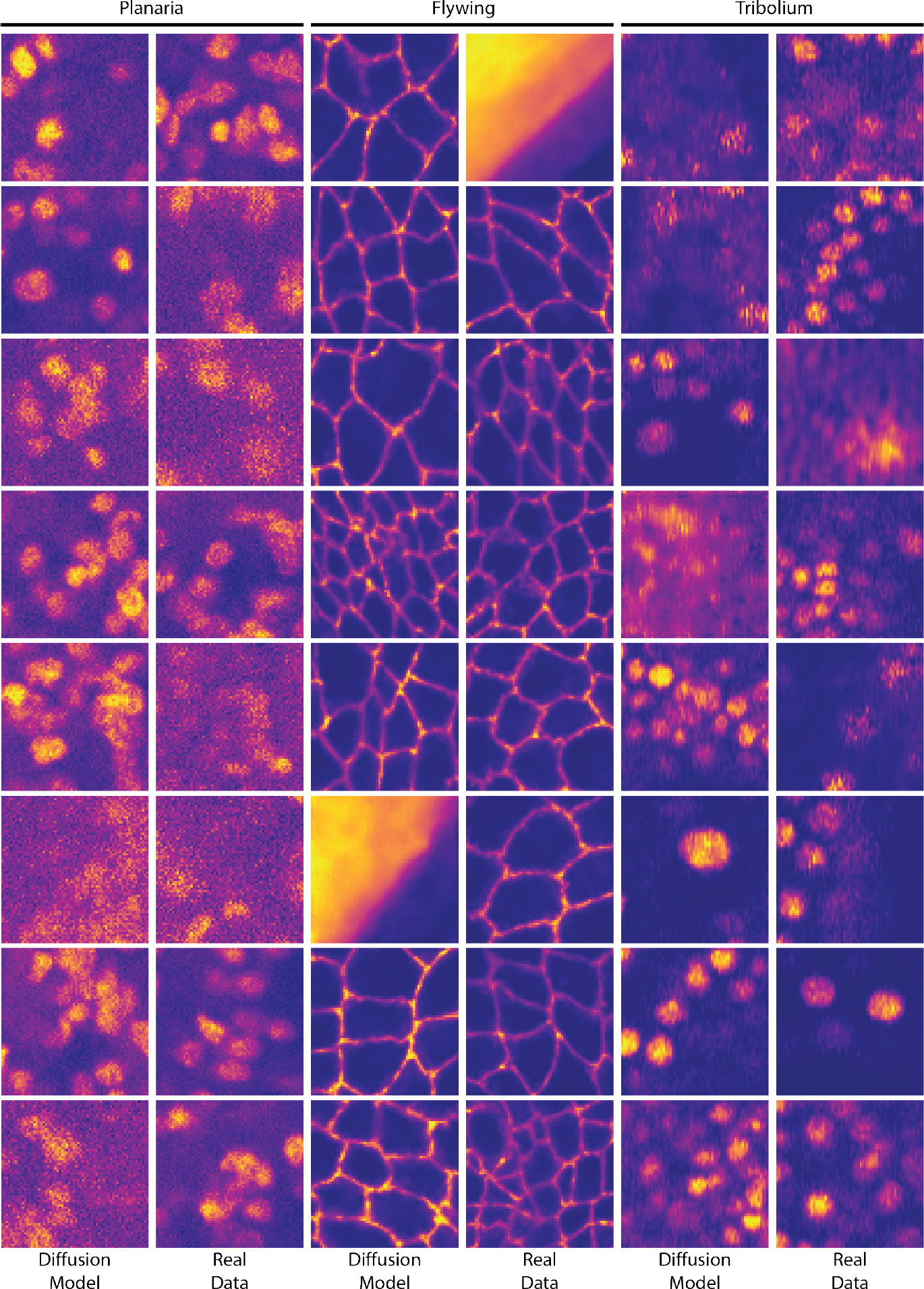
Comparison of random samples from a) diffusion model trained on clean images and b) real clean images for *Planaria*, *Flywing* and *Tribolium* datasets. Similarity of images generated by diffusion model to clean images highlight their capability to learn data distribution

Images sampled from diffusion model look perceptually similar to the clean images from corresponding datasets Fig.2 thus highlighting the superior capability of diffusion models to learn data-distribution. Such information learnt by the diffusion backbone can provide important cues during denoising in extreme noise scenarios and is the key adavantage of our proposed method.

**Table 1.**
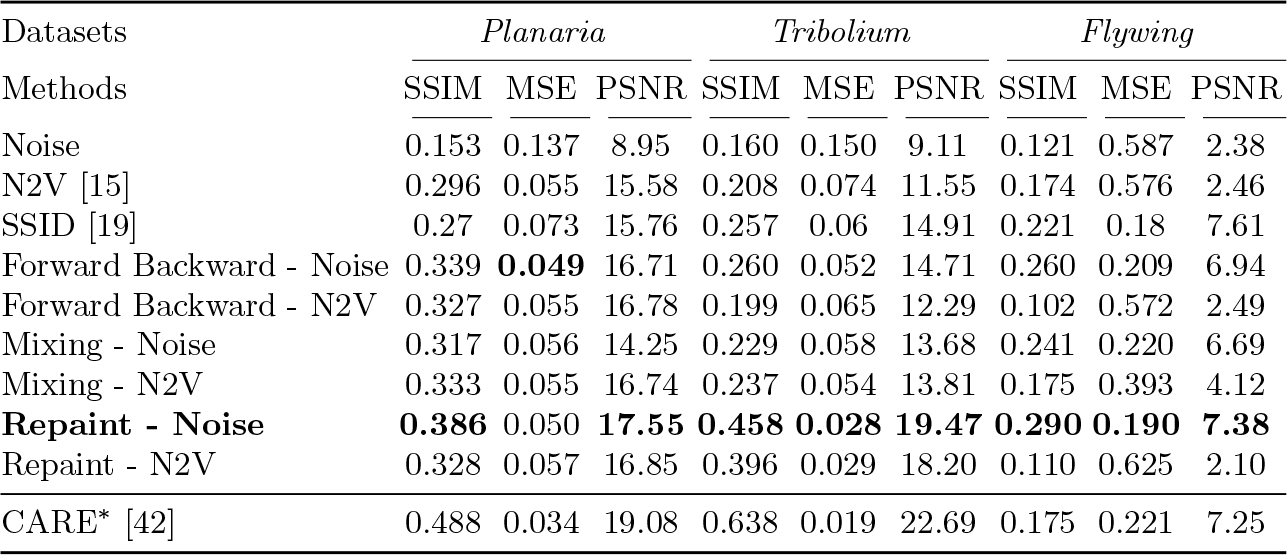
SSIM(*↑*), MSE(*↓*) and PSNR(*↑*) metrics on test sets. *^∗^*CARE is trained using paired dataset.

### 3.2 Quantitative results

We compare Baikal against state of the art methods for denoising fluorescence microscopy images. These include self-supervised method N2V [15] and paired-denoising method CARE [42]. We train both methods using author provided code and hyperparameters on the same train sets as used for training diffusion backbones. For the *Tribolium* dataset, we observed bad performance of N2V due to correlated noise in the input images and therefore trained structN2V [3] using a square mask of size 5x5 pixels around each masked pixel. For the *Flywing* dataset, since only 2D clean images were available, we trained CARE on max-projected noisy images without using a projector network as done previously [42]. We report SSIM [41], MSE and PSNR metrics to compare performance of all methods. To be consistent with CARE, we evaluate metrics on 3D datasets by first max-projecting them across the z-plane dimension.

We observe from Table. 1 that a simple conditional sampler like ForwardBackward achieves better performance than N2V across all three datasets signifying the advantage of generative priors provided by the diffusion model. Surprisingly, when N2V predictions were used as input to Forward-Backward sampler, the performance drops. We hypothesize that this could be because N2V predictions destroy underlying signal leading to sub-optimal generations. Furthermore, we observe Repaint with mixing noisy image is the best conditional sampler for all three datasets trailing only behind paired-denoising method CARE (see Fig.3). N2V required manual tuning of the masks during training to work on *Tribolium* images with correlated noise profiles. In addition, CARE required a separate projector network [42] to be trained when 3D noisy and 2D clean images in *Flywing* data are available. In contrast, our proposed approach does not require any modifications while training the diffusion models on different datasets spanning different noise profiles and projections thus making it easy to use in practical settings.

**Fig. 3.**
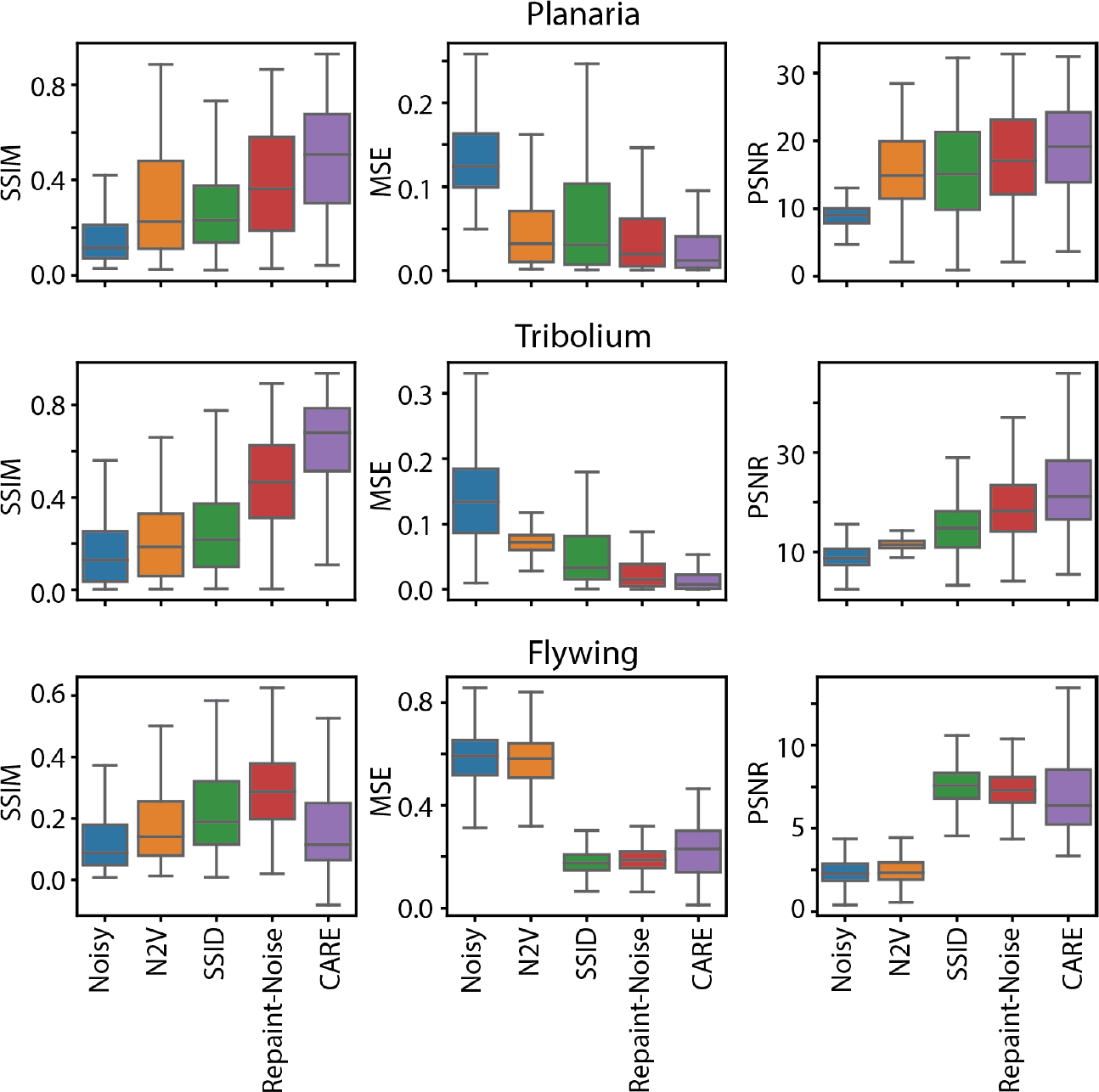
SSIM(*↑*), MSE(*↓*) and PSNR(*↑*) metrics on test sets of *Planaria* (Top), *Trbolium* (Middle) and *Flywing* (Bottom) datasets. *^∗^*CARE is trained using paired dataset.

**Fig. 4.**
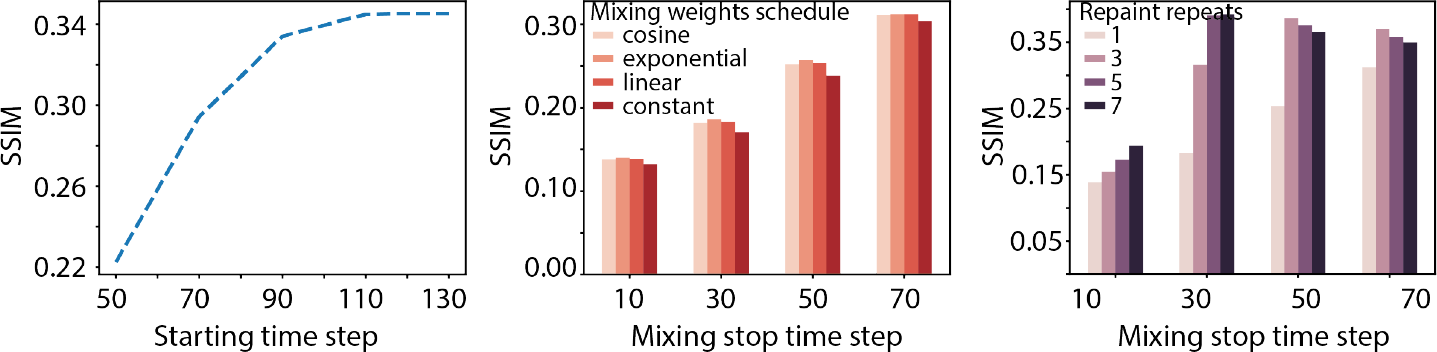
Hyperparameter effects on SSIM accuracy evaluated on *Planaria* dataset. **(Left panel)** Starting time step. **(Center panel)** Mixing stop time step and mixing weights schedule. **(Right panel)** Mixing stop time step and number of repaint repeats.

### 3.3 Ablations

To observe the sensitivity of the hyperparameters in different samplers, we vary critical parameters in each sampler and report the corresponding performance achieved on *Planaria* dataset. As seen from Fig.4 (left), increasing *t^′^* leads to cleaner generations for forward-backward sampler and the performance plateaus after 100 time steps. For mixing sampler Fig.4 (middle), mixing at time steps closer to *t* = 0 leads to decrease in performance as it re-introduces noisy pixels in the denoised output thus reverting denoising. Further, the annealing schedule for the weights minimally affects the results. In the Repaint sampler Fig.4 (right), increasing the number of repeats significantly improves the performance supporting our argument that semantic consistency is improved by repainting. Additionally best results are obtained when mixing is stopped after *t* = 50 while repainting continues.

### 3.4 Qualitative results

Qualitative comparisons highlight more advantages of our proposed framework. We observe N2V predictions contain horizontal artifacts in *Planaria* dataset (Fig.5, Supplementary Fig.1 & Supplementary Fig.4). Presence of such artifacts in N2V denoised images have been reported before [14]. Further, N2V suffers from bad performance on the *Tribolium* dataset due to the presence of correlated noise (Fig.6, Supplementary Fig.2 & Supplementary Fig.5) even after tuning the masking window size. Finally, it fails to recover cell boundaries in *Flywing* data in extremely noisy regions (Fig.7, , Supplementary Fig.3 & Supplementary Fig.6). In contrast, the best conditional sampler performs well across all scenarios. Notably in the *Flywing* dataset, Baikal is able to recover cell boundaries in extremely noisy regions (highlighted by arrows in Fig.7). In addition, while CARE predictions achieve high SSIM accuracy, qualitatively the cellular structures look extremely smooth, removing any subcellular features (Fig.5, Fig.6). In comparison, Baikal is able to preserve such subcellular features (highlighted by arrows).

**Fig. 5.**
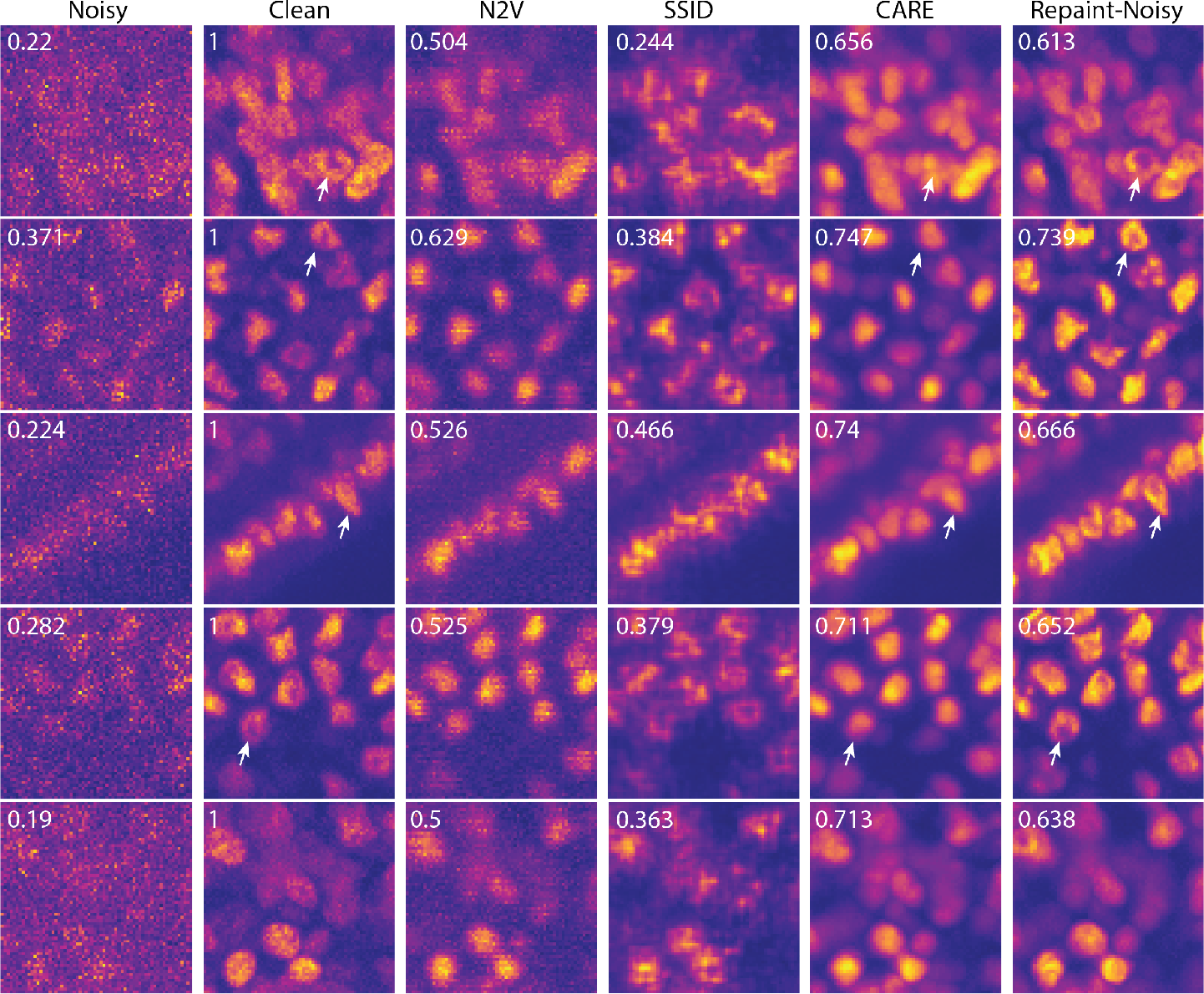
Denoising performance of methods for *Planaria* dataset. Numbers indicate SSIM w.r.t clean images. Arrows highlight examples of features preserved by our method but missed by other methods.

**Fig. 6.**
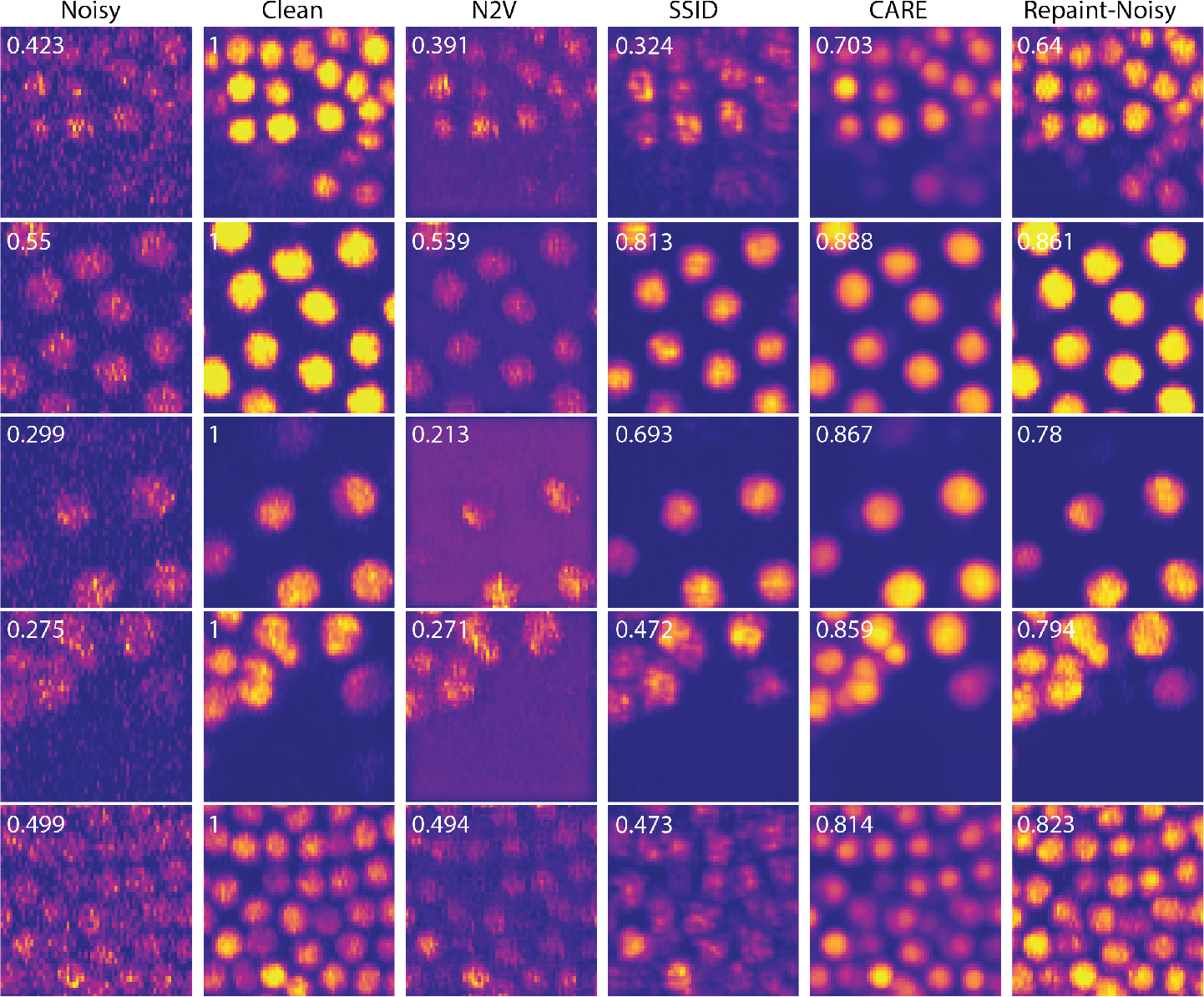
Denoising performance of methods for *Tribolium* dataset. Numbers indicate SSIM w.r.t clean images.

**Fig. 7.**
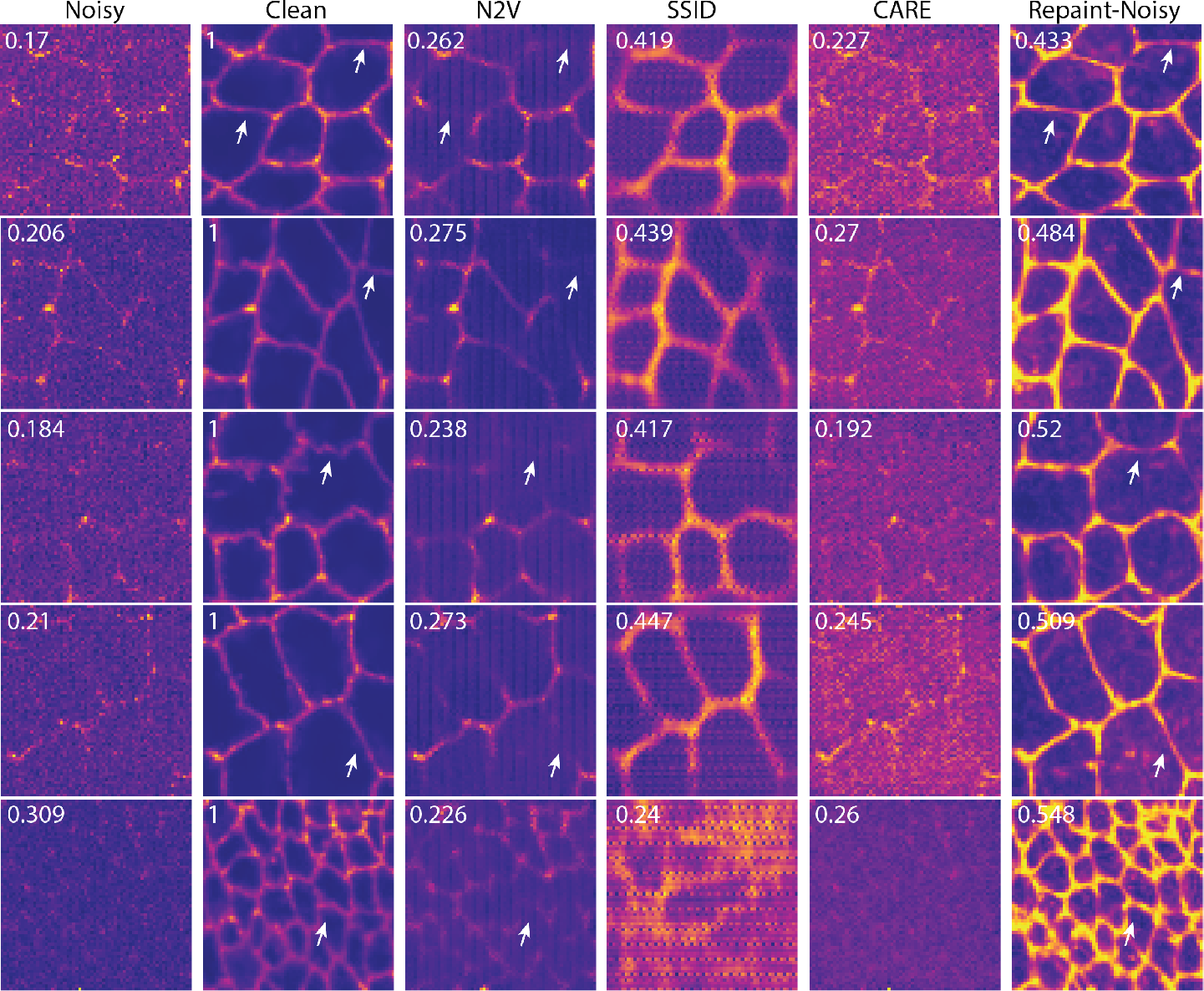
Denoising performance of methods for *Flywing* dataset. Numbers indicate SSIM w.r.t clean images. Arrows highlight examples of features preserved by our method but missed by other methods.

## 4 Conclusion

We demonstrate for the first time, to the best of our knowledge, application of Diffusion Models for denoising fluorescence microscopy images without needing paired training data. Baikal opens up future avenues for tackling other common tasks in fluorescence microscopy like de-blurring, isotropic reconstruction and super-resolution.

## 5 Supplementary Figures

**Fig. S1.**
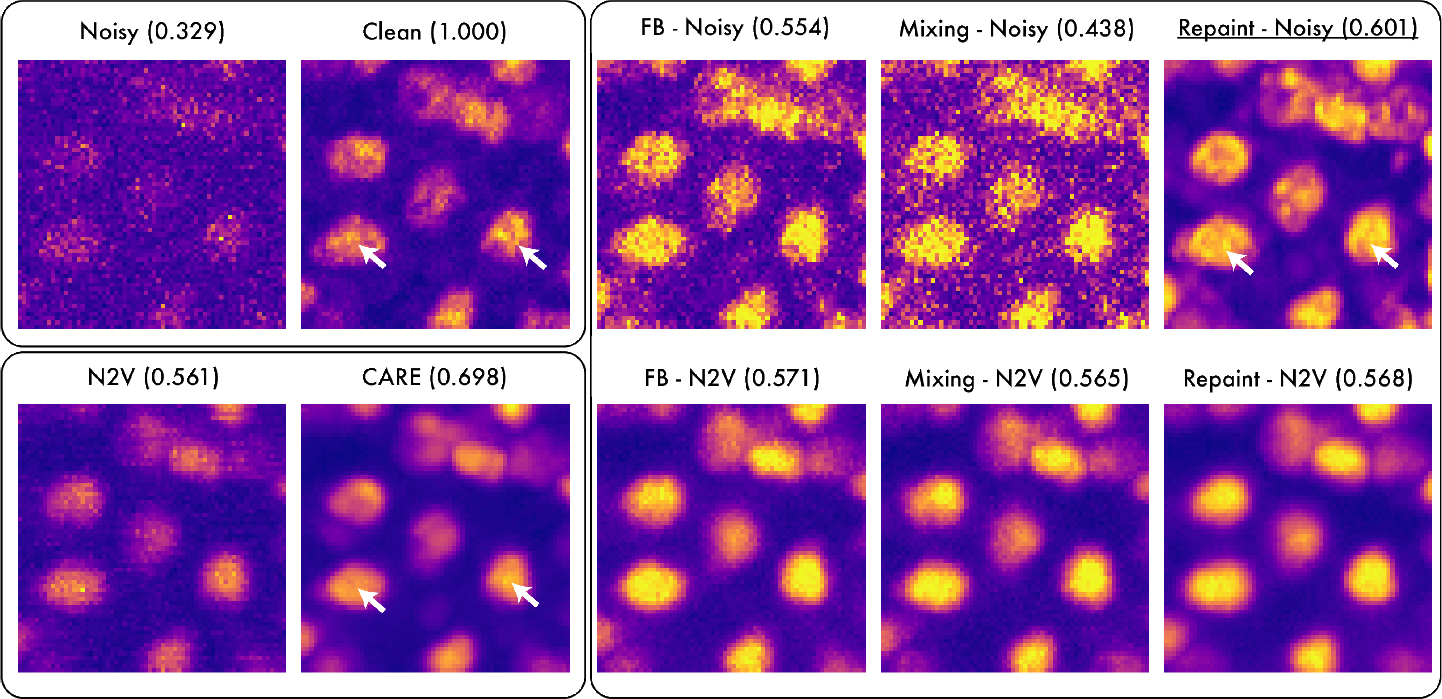
Qualitative examples comparing performance of diffusion model based denoising strategies Forward-Backward, Mixing, and Repainting. Sample predictions for *Planaria*, SSIM in brackets. **(Top left panel)** Noisy and Clean images. **(Bottom left panel)** N2V and CARE predictions. **(Right panel)** Conditional diffusion denoising using noisy image and N2V prediction as inputs.

**Fig. S2.**
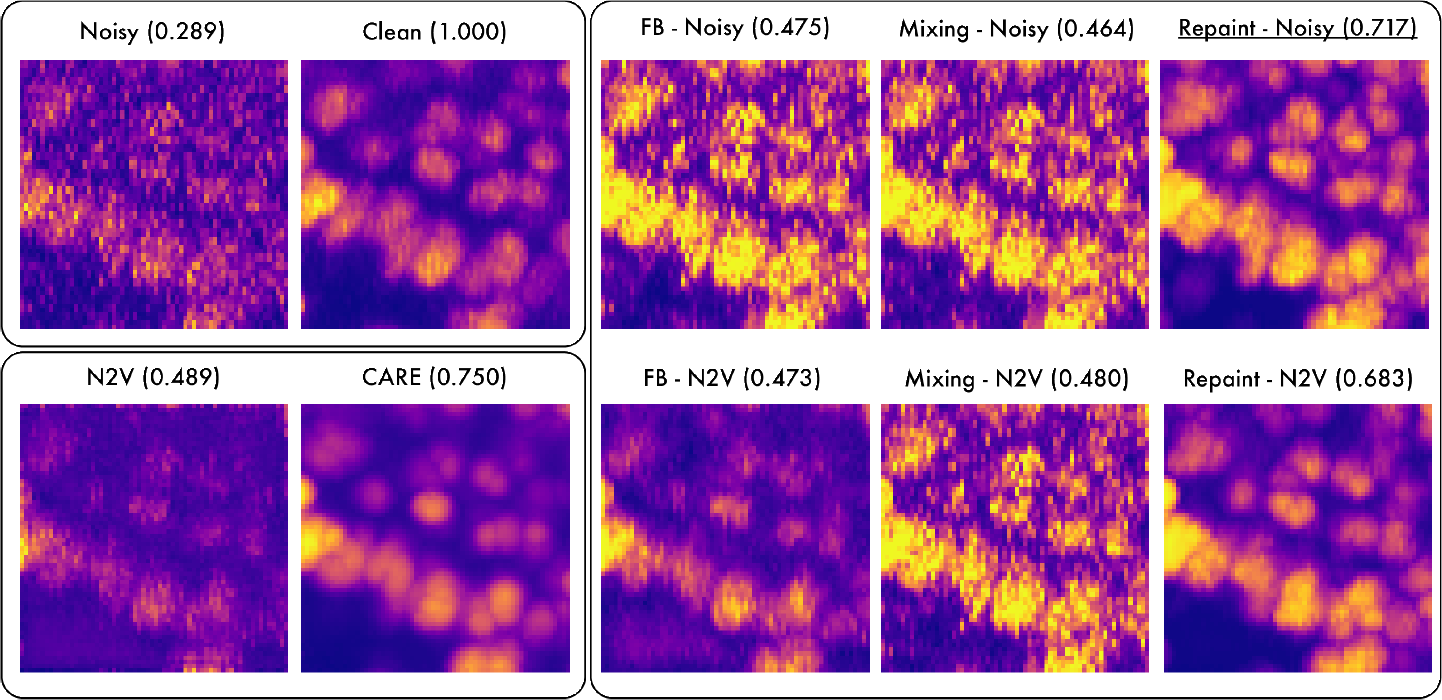
Qualitative examples comparing performance of diffusion model based denoising strategies Forward-Backward, Mixing, and Repainting. Sample predictions for *Tribolium*, SSIM in brackets. (Panels are same as Supplementary Fig.**??**)

**Fig. S3.**
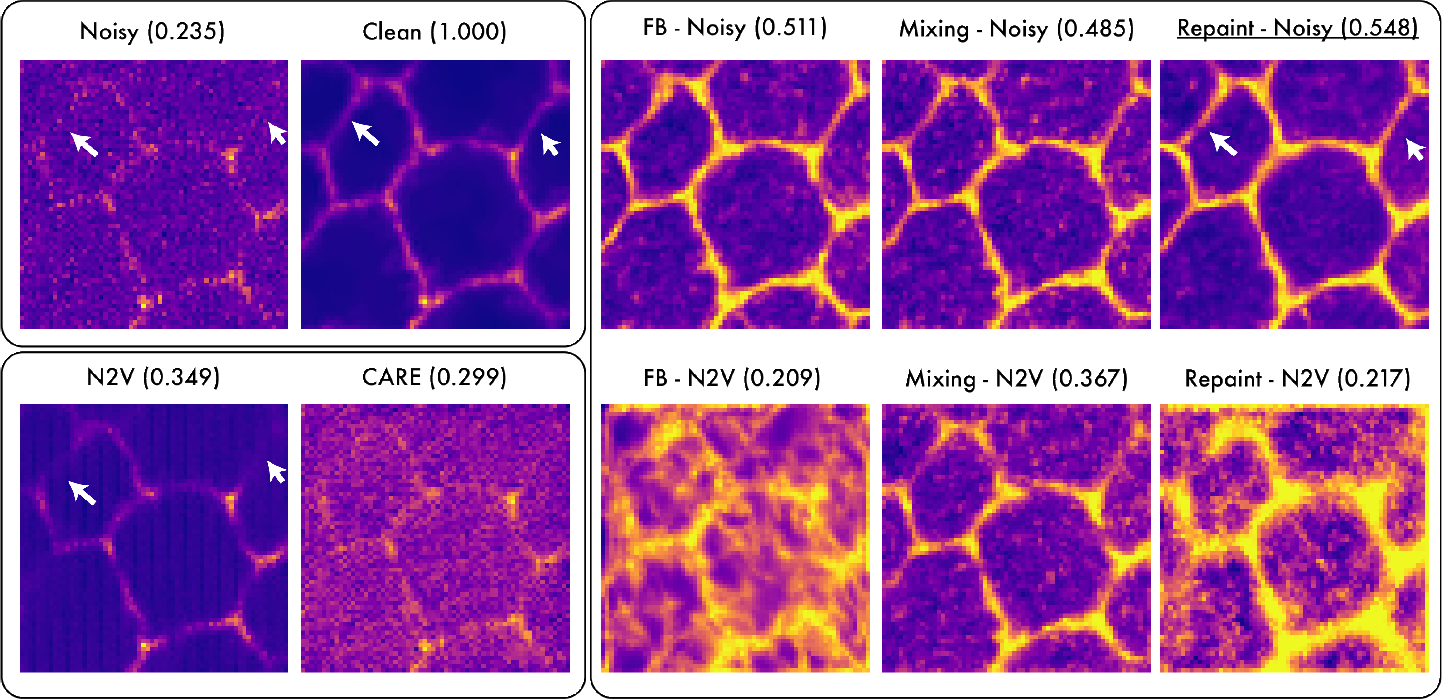
Qualitative examples comparing performance of diffusion model based denoising strategies Forward-Backward, Mixing, and Repainting. Sample predictions for *Flywing*, SSIM in brackets. (Panels are same as Supplementary Fig.**??**)

**Fig. S4.**
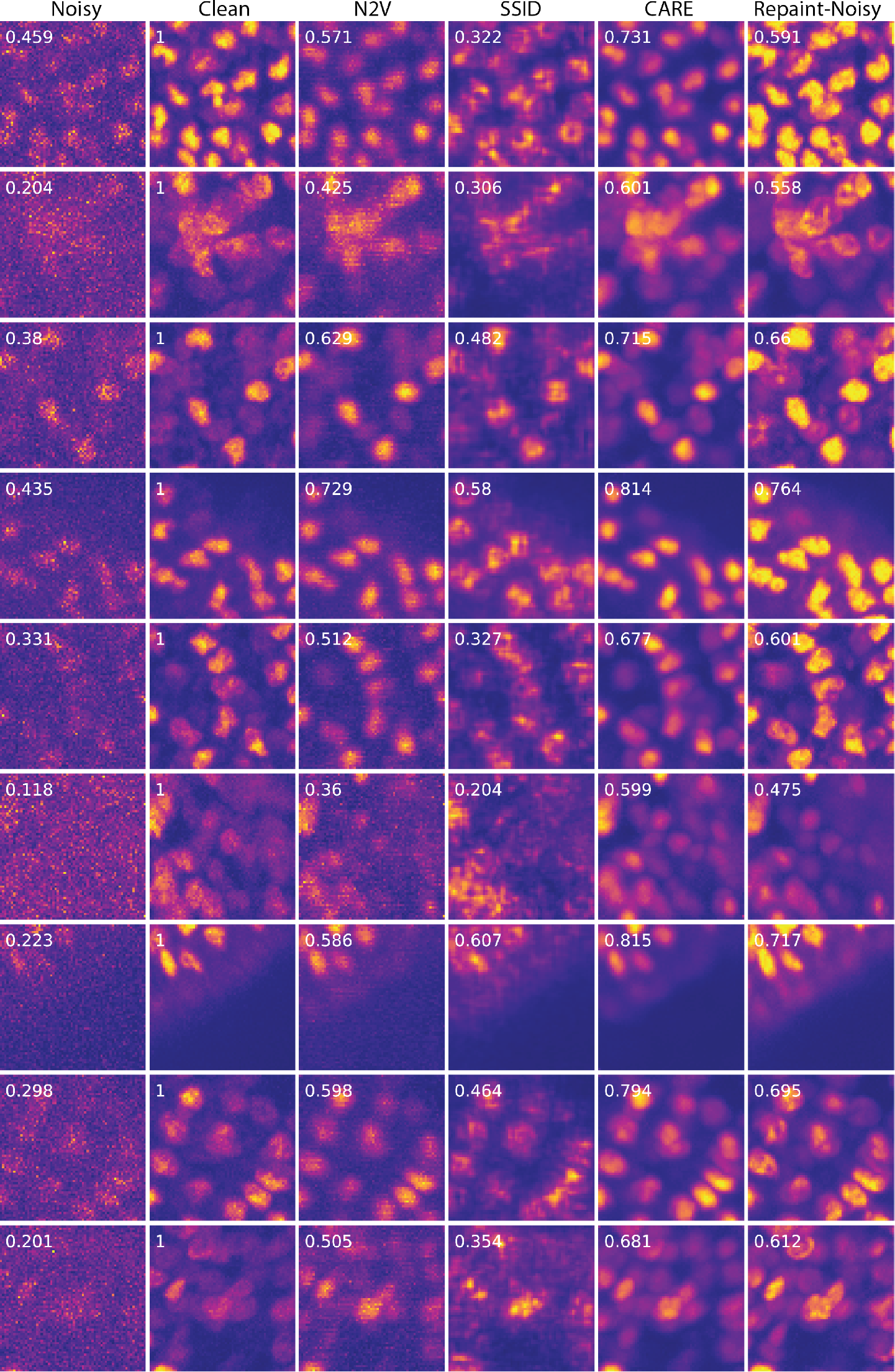
Additional examples of denoising performance of methods for *Planaria* dataset. Numbers indicate SSIM w.r.t clean images.

**Fig. S5.**
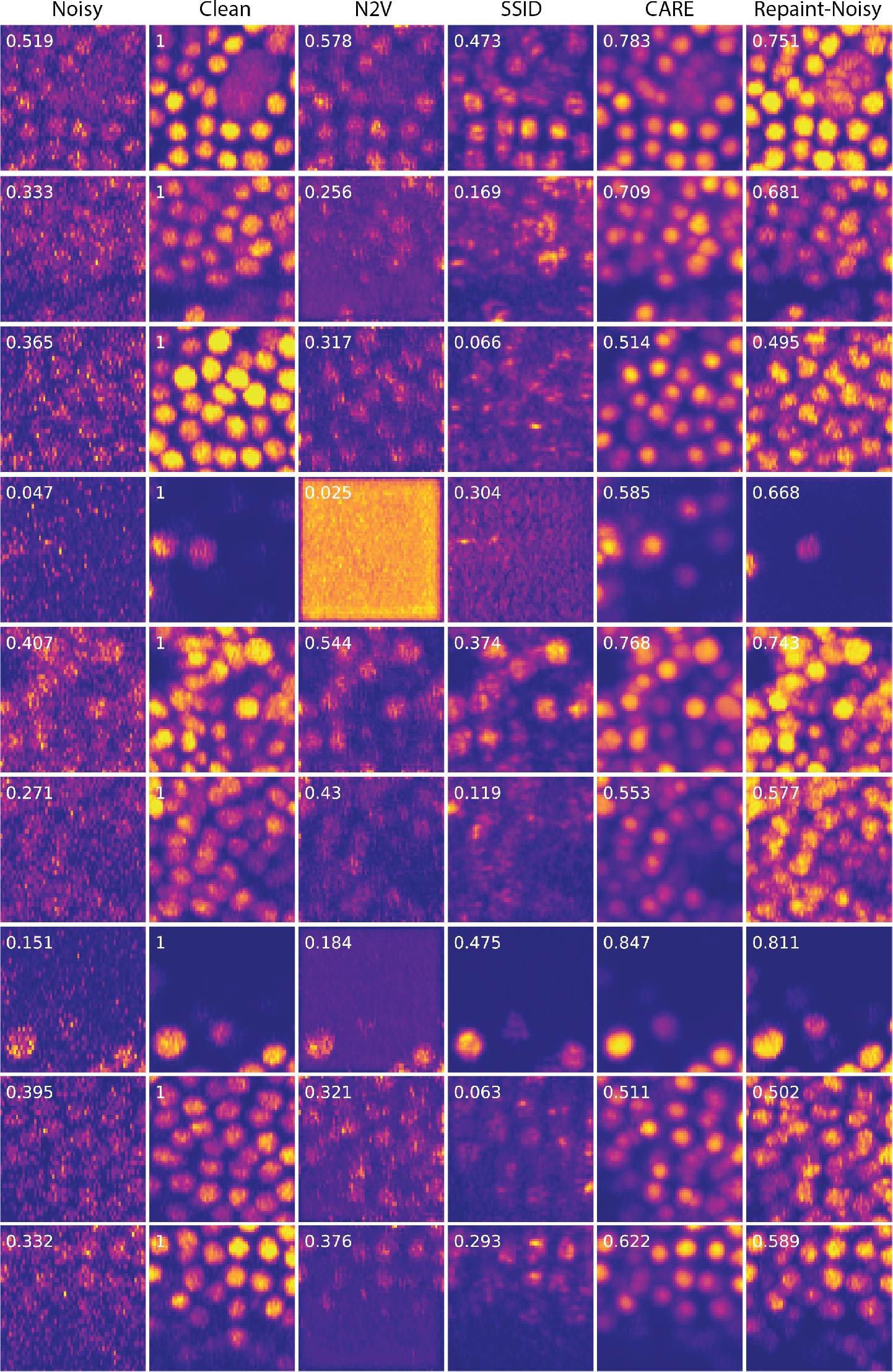
Additional examples of denoising performance of methods for *Tribolium* dataset. Numbers indicate SSIM w.r.t clean images.

**Fig. S6.**
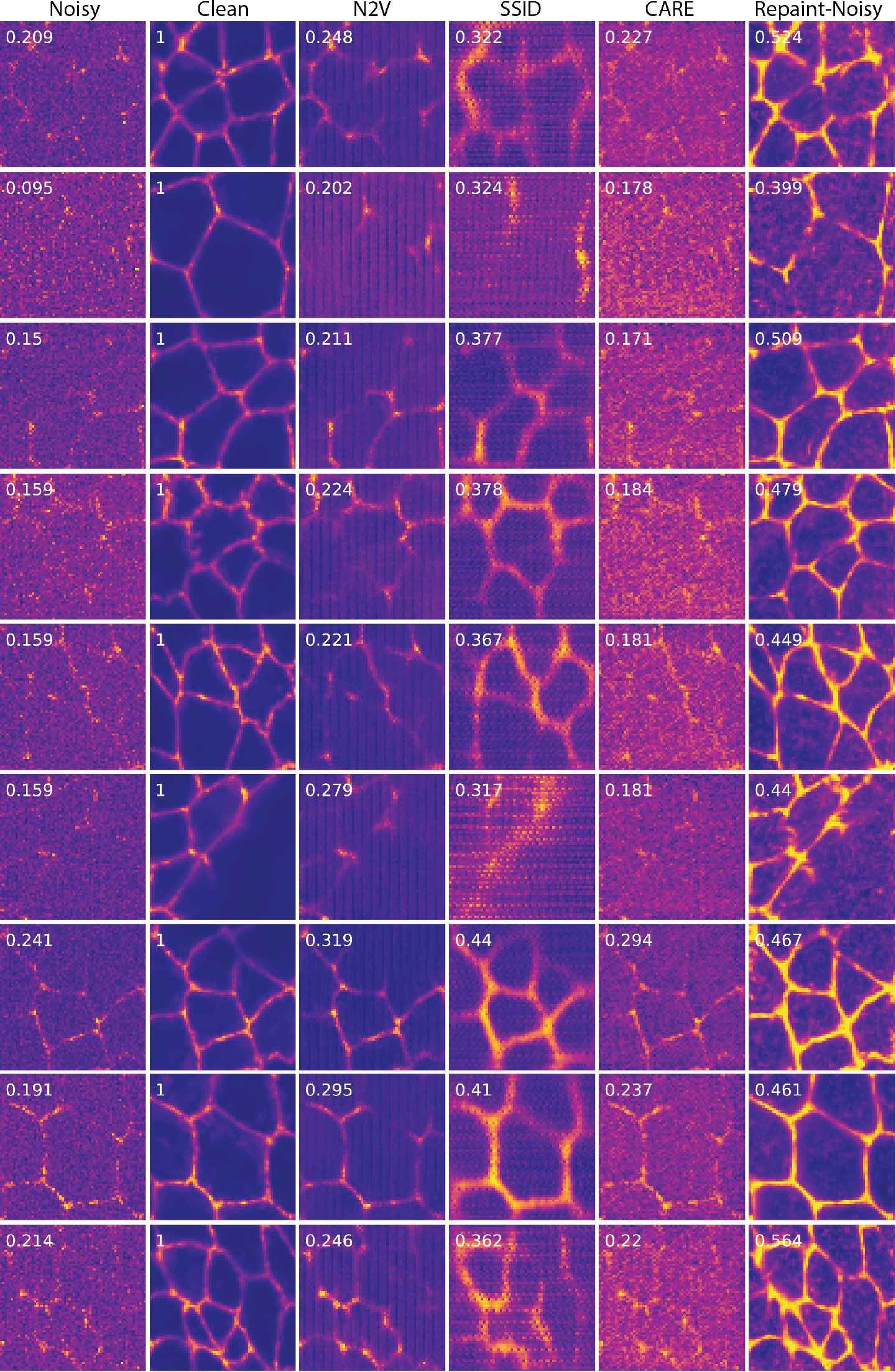
Additional examples of denoising performance of methods for *Flywing* dataset. Numbers indicate SSIM w.r.t clean images.

## Footnotes

* https://github.com/zsyOAOA/DifFace

## Notes

### Competing Interest Statement

The authors have declared no competing interest.

